# Conserved angio-immune subtypes of the cancer microenvironment predict response to immune checkpoint blockade therapy

**DOI:** 10.1101/2021.12.23.470799

**Authors:** Madhav Subramanian, Ashraf Ul Kabir, Derek A.G. Barisas, Karen Krchma, Kyunghee Choi

**Author notes:** Corresponding Author, Dr. Kyunghee Choi.

## Abstract

Tumor microenvironment (TME) shapes the tumor progression and therapy outcome. Particularly, tumor angiogenesis and immunity impact the effect of immune checkpoint blockade (ICB) therapy. Here, we analyzed the transcriptome from 11,069 patients from The Cancer Genome Atlas (TCGA) to assess 91 functional gene sets corresponding to endothelial and T-cell activity. Intriguingly, TME across 30 non-hematological tumors can be classified into three distinct conserved angio-immune subtypes: high angiogenesis with low immune activity, low angiogenesis with high immune activity, and the one in-between. Remarkably, patients displaying TME with poor angiogenic activity with corresponding high immune activity show the most significant responses to ICB therapy in many cancer types. Notably, re-evaluation of the Javelin Renal 101, renal cell carcinoma clinical trial, provided compelling evidence that the baseline angiogenic state is critical in determining responses to checkpoint blockade. This study offers a clear rationale for incorporating baseline angiogenic state for ICB treatment decision-making.

---

Heterogeneity in the tumor cells and the surrounding microenvironment drives treatment resistance among patients ^1^. As such, there have been efforts to identify hierarchical classifications of tumor subtypes on a pan-cancer level ^2, 3, 4, 5, 6^. Unlike the genetically unstable tumor cells, microenvironmental features are more stable and could be conserved across tumor types. As such, delineating tumor microenvironment (TME) subtypes may enable the identification of common resistance mechanisms across tumor types and guide treatment decision-making, particularly for treatments targeting microenvironmental features such as anti-angiogenic agents and immune checkpoint blockade (ICB).

Abnormal tumor vasculature contributes to tumor growth and escape by remodeling the local TME where tumors thrive. Recent evidence from pre-clinical studies suggests that vascular dysfunction in tumors presents physical and chemical barriers to the infiltration of immune cells. Hyperpermeable, immature blood vessels result in improper circulation and hypoxia in the TME and cannot adequately perform as a conduit for cytotoxic T-cell trafficking to the TME ^7^. Moreover, under sub-optimal perfusion, glycolytic co-option produces a highly acidic environment that suppresses T-cell effector functions ^8^. Additionally, angiogenic tumor endothelial cells express death signals like FAS ligand (FASLG) that induce apoptosis in infiltrating cytotoxic T-cells ^9^. Importantly, recent pre-clinical evidence suggests that normalization of vasculature can reverse the immunosuppressive milieu, promotes anti-tumor T-cell immunity, and synergizes with ICB, suggesting a close interaction that is beyond mere correlative ^10, 11, 12^.

While ICB has revolutionized the treatment of metastatic and solid tumors by providing lasting regression to a subset of patients, a sizable majority of patients fail to mount responses to ICB ^13^. High interdependence of angiogenesis and T-cell infiltration and activation in the TME raises the question whether the baseline angiogenic state can be a determinant of tumor immune microenvironment characteristics and can be prognostic of ICB response in patients. Elucidating heterogeneity in the baseline angiogenic state and corresponding immune activation might help inform treatment decision-making, allowing us to reduce treatment-related toxicities faced by patients and design more efficient therapeutic regimens.

Here, we show that the baseline angiogenic state is a determinant of cytotoxic T-cell infiltration and function in the TME. We developed a transcriptional profile analysis pipeline to stratify patients based on baseline angiogenic and immune activity using endothelial cell and T-cell functional genesets. This highly interpretable tool provides insight into the intimate interaction between angiogenic and immune processes and enables us to identify pan-cancer molecular subtypes of tumors. Molecular subtypes based on baseline angiogenic and T-cell activity are prognostic of response and survival of patients treated with ICB. Retrospective analysis of pre-treatment datasets reveals that highly angiogenic patients do not derive clinical benefit from FDA-approved first-line ICB strategies. We present a reformed understanding of how angiogenesis and T-cell immunity are linked across tumor types, laying the foundation for more efficient treatment decision-making approaches.

## RESULTS

### Angiogenesis and T-cell mediated Immunity are Inversely Correlated Across Cancer Types

To characterize the interplay between angiogenesis and T-cell mediated immunity in tumors, we deconvoluted the TME’s angiogenic and T-cell functional landscape from all solid tumor samples from The Cancer Genome Atlas (TCGA). Transcriptomic data from 30 non-hematological tumor types were used to score 91 functional gene sets corresponding to endothelial and T-cell activity from the Molecular Signatures Database (MSigDB) using Gene Set Variation Analysis (GSVA, Fig. 1a, Table S1). Endothelial cell activity gene sets included gene sets relating to angiogenesis, endothelial cell chemotaxis, and endothelial cell signaling downstream of angiocrine factors. T cell activity gene sets included gene sets relating to T cell activation, T cell differentiation, T cell cytotoxicity, and tissue infiltration of T cells.

**Figure 1.**
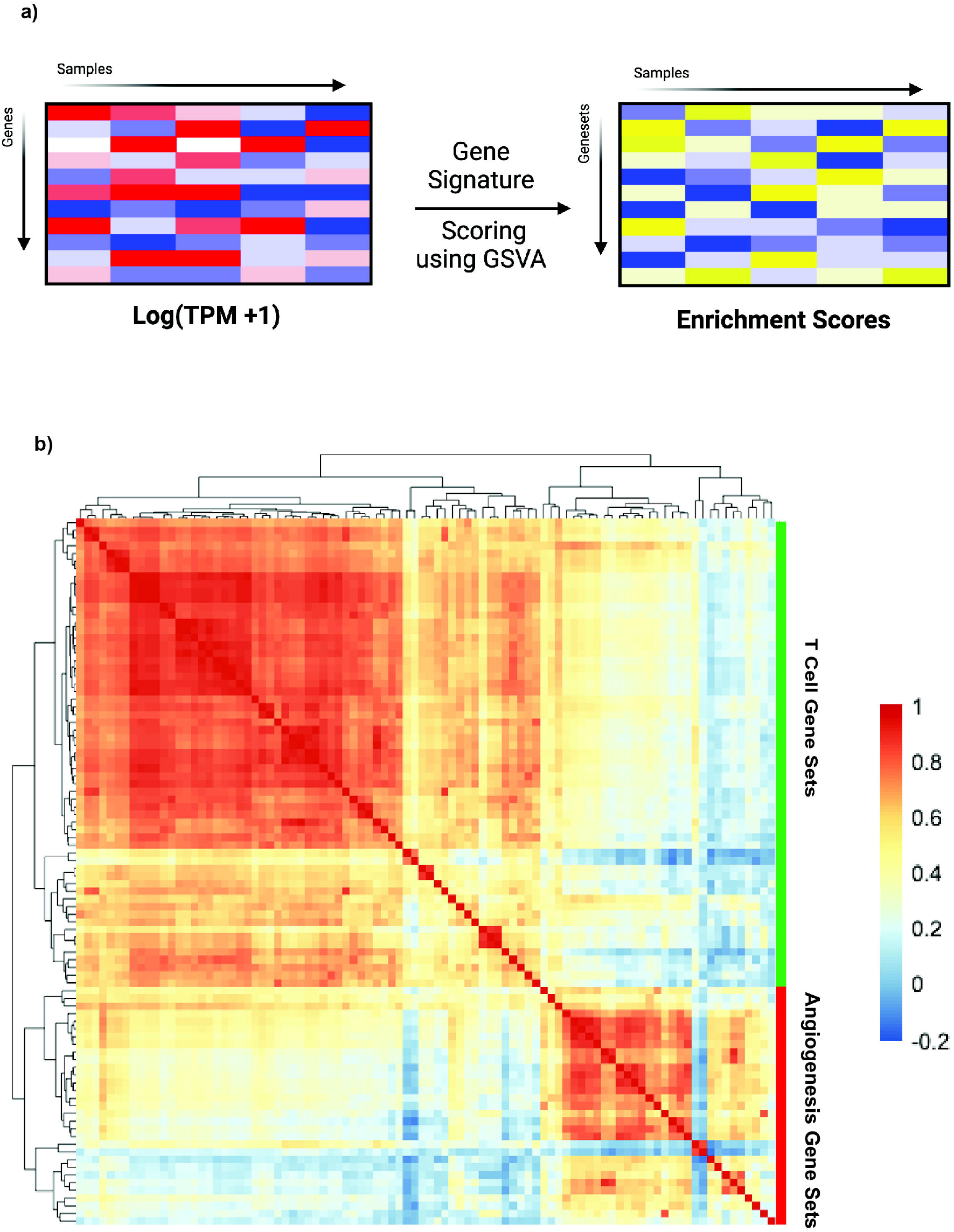
Characterizing gene set relationships. a) Schematic depicting generation of the enrichment matrices used for the angio-immune subtype identification across tumor types. b) Heatmap depicting Pearson correlation of angiogenesis and T-cell gene sets across 11,069 TCGA tumor samples.

We first investigated the distributions of gene sets across patients harboring non-hematological tumor types. A correlation matrix generated by using Pearson Correlation Coefficients (PCC) was built that revealed two functional gene set modules bound by positive correlation (Fig. 1b). Gene sets corresponding to angiogenic function including, endothelial cell migration, proliferation, sprouting, and angiogenesis, were the characteristics of one module. The other module was characterized by gene sets corresponding to T-cell effector function, antigen processing and presentation, and trans-endothelial migration of leukocytes. We found that T cell gene set scores were inversely correlated with angiogenic gene set scores (Fig. 1b). This inverse relationship between tumor angiogenesis and immunity has also been observed in mouse models in a previous study^11^.

### Identification of The Angio-Immune Subtypes

A distinct inverse relationship between angiogenic and immune gene sets suggests that we may be able to stratify patients based on baseline angiogenic and T-cell activity. To identify the angio-immune subtypes of TME based on the distinct distribution of signatures associated with angiogenic and T-cell functions, we generated a correlation matrix of patients using the distinct distribution of gene signature enrichment across samples. As a result, distinct subtypes of the TME were uncovered and termed as C1, C2, and C3 (Fig. 2a). With the challenge of current standard of care, patients from the three angio-immune subtypes had negligible differences in survival (Fig. 2b). These subtypes were conserved across all tumor types; however, the proportion of the angio-immune subtypes varied across tumor types (Fig. 2c). Further analyses revealed that distinct distribution of gene sets characterized the angio-immune subtypes. As such from the angiogenesis perspective, C1 contained strong positive enrichment for angiogenic gene sets, including endothelial cell (EC) migration, EC proliferation, EC sprouting. In contrast, C3 was characterized by the downregulation of functional angiogenesis signatures. C2 had no significant enrichment for functional angiogenesis signatures (Fig. 2d). From the T-cell immunity perspective, C1 had marked downregulation of T-cell functional signatures, including T-cell chemotaxis, T-cell mediated cytotoxicity, T-cell extravasation, and antigen presentation to T-cells. Conversely, C3 presented with strong positive enrichment for T-cell functions. Again, C2 had no significant enrichment for T-cell-related gene sets either. This analysis provides an easily interpretable framework to classify TMEs independent of the anatomical location of the tumor.

**Figure 2.**
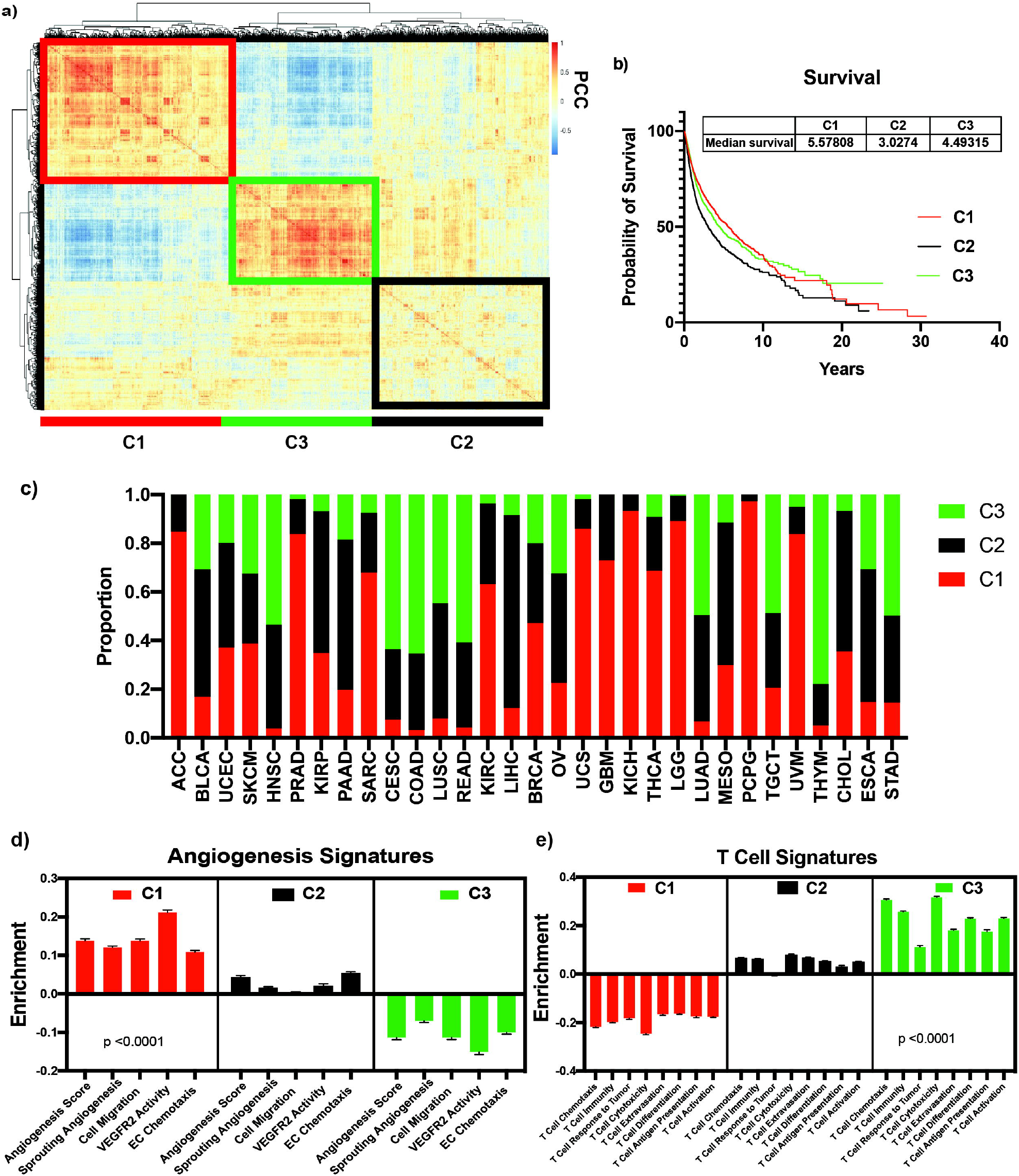
Three pan cancer angio-immune subtypes identified. a) Heatmap of Pearson Correlation of patients with non-hematological tumor types across 91 gene sets corresponding to T-cell and angiogenesis activity b) Overall survival of patients belonging to the 3 angio-immune subtypes across tumors. c) Stacked barplots depicting relative proportion of each angio-immune subtype in all cancers queried d) Bar graphs depicting the average enrichment of angiogenesis signatures in the three angio-immune subtypes. e) Bar graphs depicting the average enrichment of T-cell signatures in the three angio-immune subtypes.

### Immune Characteristics of The Angio-Immune Subtypes

To dissect the tumor immune microenvironment of the angio-immune subtypes, a signature dependent method to estimate enrichment of cell types in the TME, xCell^14^, was used to identify the enrichment of 64 different immune and stromal cell types across the angio-immune subtypes. Notably, there was a more significant enrichment of dendritic cells, CD8+ T-cells, B cells, Th1 and Th2 cells, and M1 macrophages in C3 (Fig. 3a). Conversely, C1 had a distinct increase in enrichment of endothelial and fibroblast cells and poor enrichment for immune cell subtypes (Fig. 3a, b). C1 may contain a denser stroma, evidenced by a higher stromal score (Fig. 3b). Infiltrating CD8+ T-cells in C3 may have a more significant effector function as evidenced by increased expression of co-stimulatory molecules in C3 compared to C1 and C2 (Fig. 3c). Infiltrating CD8 T-cells also display activation as evidenced by the upregulation of exhaustion markers (Fig. 3d).

**Figure 3.**
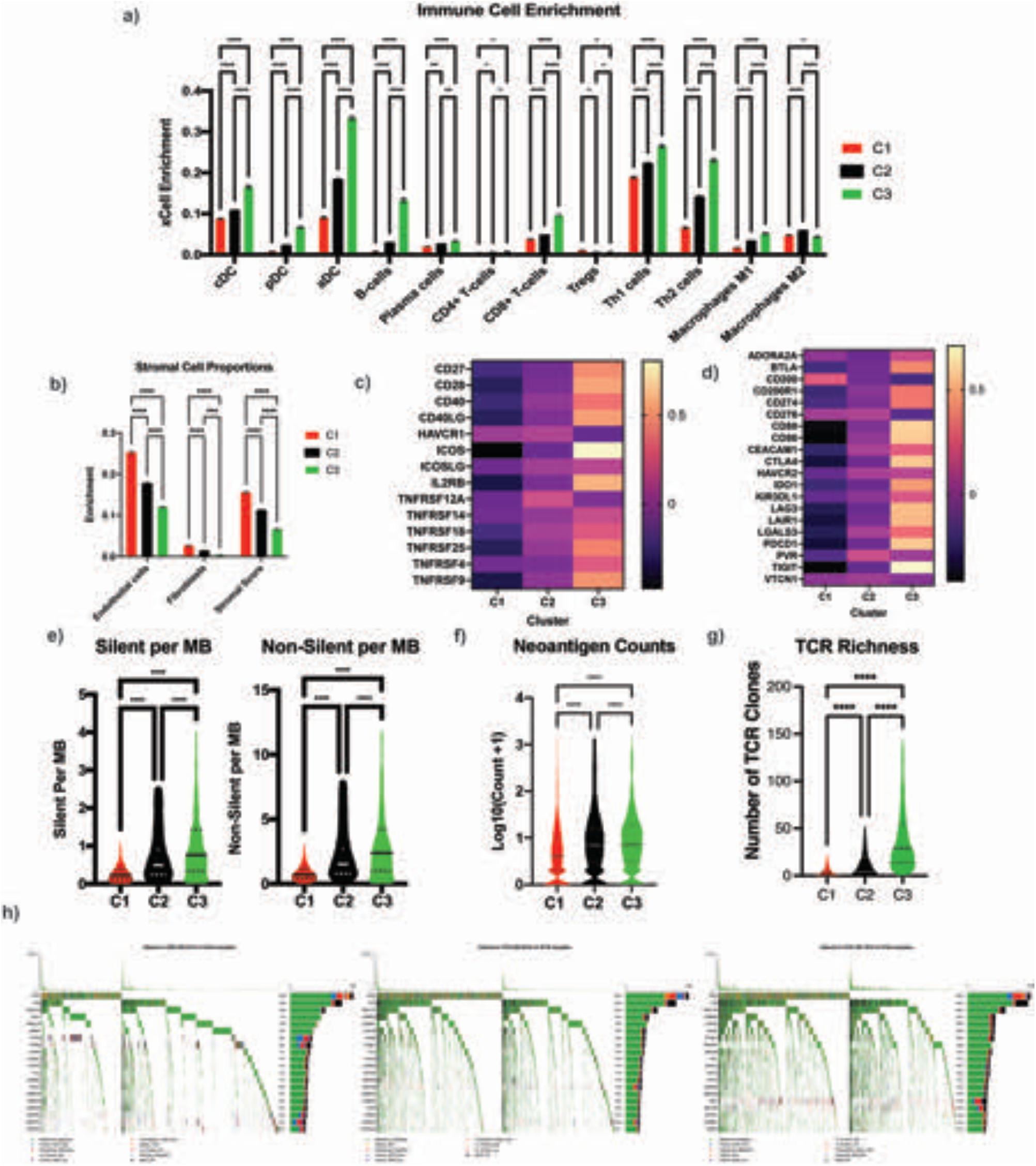
Immune characteristics of angio-immune subtypes. a) Bar plots showing xCell enrichment results for major tumor hematopoietic cells across angio-immune subtypes b) Bar plots showing xCell enrichment results for endothelial cells, fibroblasts, and pericytes across angio-immune subtypes c) Heatmap of z-scored expression co-stimulatory molecules in three angio-immune clusters d) Heatmap of z-scored expression exhaustion/activation markers in three angio-immune clusters e) Violin plots depicting silent and non-silent mutational burden across three angio-immune clusters. f) Violin plot of neoantigen counts across three angio-immune clusters g) Violin plot of TCR richness across three angio-immune clusters h) Oncoplot for overview of mutations in C1, C2 and C3 generated using maftools * *p<0*.*05*, ** *p < 0*.*01*, *** *p < 0*.*001*, **** *p < 0*.*0001*; One way ANOVA was used for comparison across more than two groups; error bars indicate SEM.

Somatic variation in tumor cells can dictate the robustness of immune responses in the TME^15^. A high tumor mutational burden in tumors is suspected of promoting the formation of antigenic peptides that can trigger immune responses. We analyzed the publicly available MC3 repository for somatic variants derived from whole-exome sequencing data to characterize somatic variations of tumors in the angio-immune subtypes. The silent and non-silent mutational burden was higher in C3 than in C2 and C1 (Fig. 3e). In line with this finding, the mean neoantigen load was greater in C3 than C2 and C1 (Fig. 3f). Infiltrating T-cells also contained greater clonality in TCR in C3 when compared to C2 and C1 (Fig. 3g). Moreover, distinct actionable mutations characterized different angio-immune subtypes. IDH1, BRAF, PTEN, and ATRX mutations were enriched within C1. CSMD3, LRP1B, ZFHX4, and USH2A mutations were enriched in C2 and C3. FAT4 mutations were enriched in C3 (Fig. 3h). The distinct immune environment of C3 showing higher CD8 T-cell infiltration, increased expression of exhaustion markers, and increased mutational burden suggests that this group of patients may benefit more from ICB.

### Response and Survival Post-ICB

Several clinical and pre-clinical studies suggest that angiogenic control heightens response to ICB ^10, 11, 12^. Additionally, there have been numerous studies suggesting that the immune profile observed in C3 (high TMB, enriched neoantigen count, improved infiltration of CD8+ T-cells, higher PD-L1 expression) corresponds to improved response to ICB ^16, 17^. As such, to evaluate if the angio-immune subtypes are prognostic of survival and response upon treatment with ICB, RNA sequencing data from pre-treatment biopsies were queried. Three cohorts of patients were assembled consisting of three different tumor types treated with ICB: metastatic melanoma, metastatic gastric cancer, and metastatic bladder cancer ^18, 19, 20, 21, 22^.

The angio-immune subtypes were largely conserved in the metastatic melanoma (Fig. 4a), metastatic gastric cancer (Fig. 4b), and metastatic bladder cancer (Fig. 4c) cohorts. The distribution of enrichment scores was also consistent with the pan-cancer analysis for metastatic melanoma (Fig. 4d), metastatic gastric cancer (Fig. 4e), and metastatic bladder cancer (Fig. 4f) cohorts. Remarkably, C3 displayed a marked increase in response rate to anti-PD1 therapy in the metastatic melanoma cohort. Improved response rate translated to an improved progression-free survival (PFS; *p = 0*.*0093*) and overall survival (OS; *p = 0*.*0027*) for patients in C3 (Fig. 4g). Melanoma patients in C3 had not reached median PFS. In comparison, the median PFS for C1 was 4.1 months, and the median PFS for C2 was 15.7 months. Similarly, metastatic gastric cancer patients treated with anti-PD1 belonging to C3 displayed a drastic improvement in response rate in comparison to C1 and C2 (Fig. 4h). Likewise, bladder cancer patients belonging to C3 displayed improved PFS than C2 and C1 (*p = 0*.*0055*; Fig. 4i). The results above clearly demonstrates that the angio-immune subtypes can be prognostic of response and survival for patients treated with ICB.

**Figure 4.**
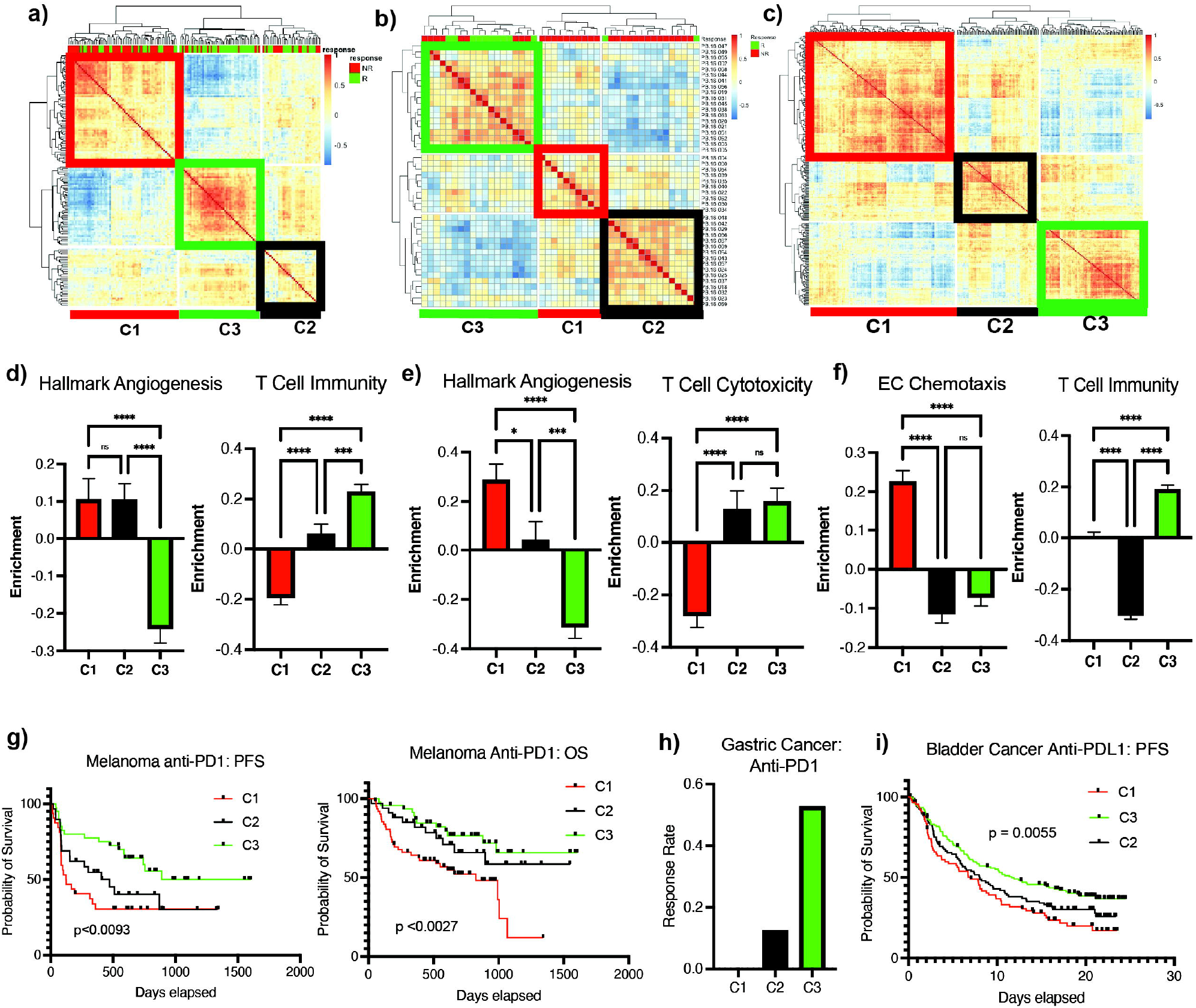
Angio-immune subtypes are prognostic of response to anti-PD1/PDL1. a) Heatmap of Pearson Correlation of 145 patients with metastatic melanoma across 91 gene sets corresponding to T-cell and angiogenesis activity. Response status is depicted for each patient as response (R = green) and non-response (NR = red). b) Heatmap of Pearson Correlation of 45 patients with metastatic gastric cancer across 91 gene sets corresponding to T-cell and angiogenesis activity. Response status is depicted for each patient as response (R = green) and non-response (NR = red). c) Heatmap of Pearson Correlation of 348 patients with metastatic bladder cancer across 91 gene sets corresponding to T-cell and angiogenesis activity. d) Bar graphs depicting the average enrichment of angiogenesis signature and T-cell signature in the three angio-immune subtypes in metastatic melanoma cohort e) Bar graphs depicting the average enrichment of angiogenesis signature and T-cell signature in the three angio-immune subtypes in metastatic gastric cancer cohort f) Bar graphs depicting the average enrichment of angiogenesis signature and T-cell signature in the three angio-immune subtypes in metastatic bladder cancer cohort g) Progression free survival (PFS) and overall survival (OS) for patients in different angio-immune clusters upon treatment with anti-PD1 in metastatic melanoma patients h) Response rate for patients in different angio-immune clusters upon treatment with anti-PD1 in metastatic gastric cancer patients. i) Progression free survival (PFS) for patients in different angio-immune clusters upon treatment with anti-PDL1 in metastatic bladder cancer patients * = p<0.05, ** = p < 0.01, *** = p < 0.001, **** = p < 0.0001; One way ANOVA was used for comparison across more than two groups; Log rank tests were used for all survival analysis; error bars indicate SEM.

### Re-evaluation of Javelin Renal 101 Clinical Trial

The approval of the combination of avelumab, a monoclonal anti-PDL1 antibody, and axitinib, a small molecule tyrosine kinase inhibitor, for the first-line treatment of metastatic renal cancer was provided in 2020 by the FDA. The approval was granted on the back of the median PFS improvement from 8.4 months with the previous standard of care, sunitinib, a tyrosine kinase inhibitor, to 13.8 months for the combination ^23^. We sought to identify if stratification by the angio-immune subtypes can confer more significant differences in response and survival.

Here, we scored the enrichment of the 91 angio-immune signatures using RNA sequencing data from pre-treatment biopsies from patients enrolled in the Javelin Renal 101 trial. The three angio-immune subtypes were conserved in this dataset (Fig. 5a). Angiogenic and immune gene sets’ enrichment distribution was consistent with the earlier observations made from analysis of TCGA data (Fig. 2d, e & Fig. 5b). C1 presented an upregulation of angiogenesis functions and a downregulation of T-cell functions. Conversely, C3 presented an upregulation of T-cell functions and a downregulation of angiogenesis functions. Unsurprisingly, C2 displayed no significant enrichment in either functional module.

**Figure 5.**
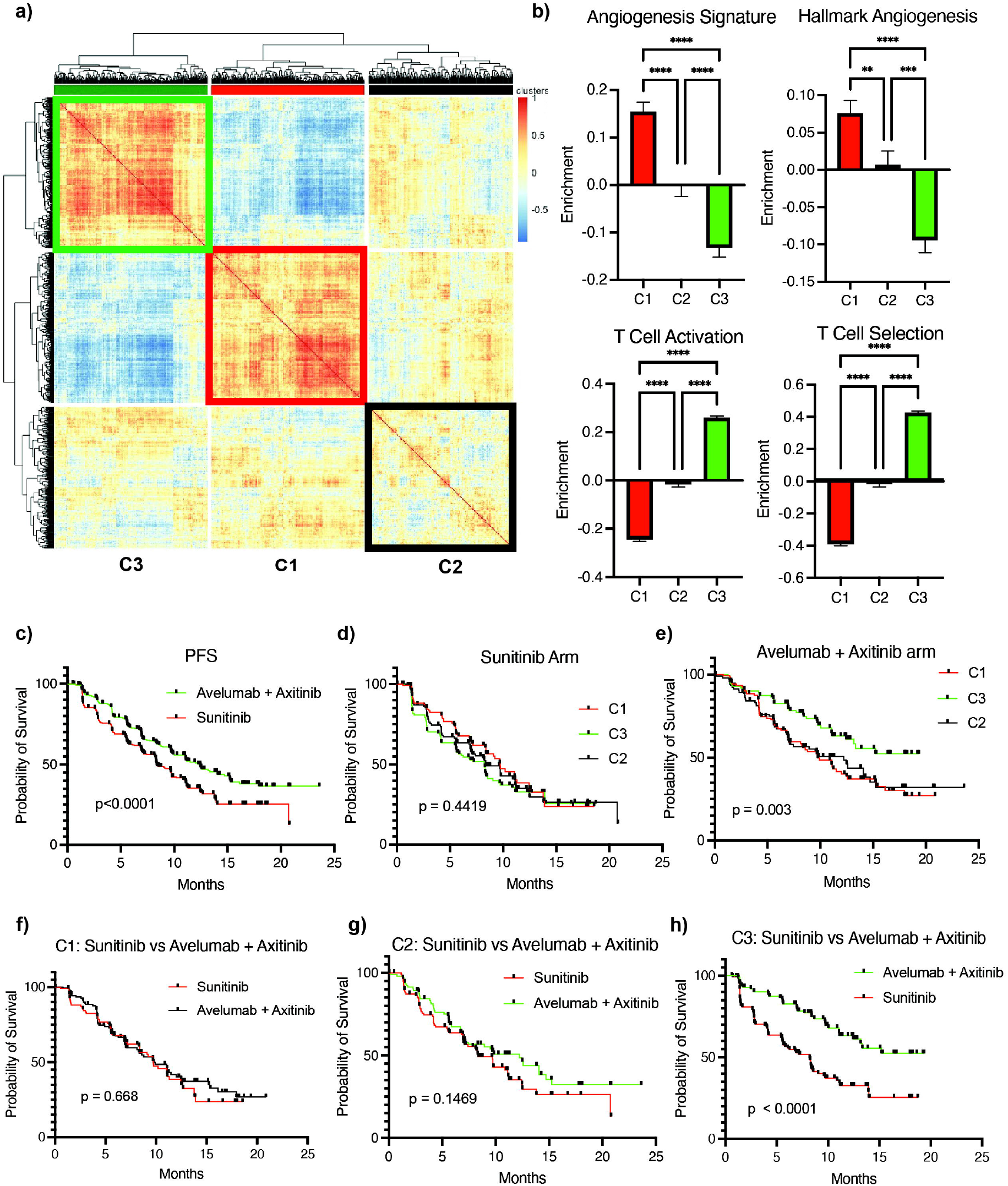
Re-evaluation of Javelin Renal 101 reveals differing propensity for response by the angio-immune subtypes. a) Heatmap of Pearson Correlation of 726 patients with metastatic renal cell carcinoma across 91 gene sets corresponding to T-cell and angiogenesis activity. b) Bar graphs depicting the average enrichment of angiogenesis signatures and T-cell signatures in the three angio-immune subtypes in metastatic renal cell carcinoma cohort c) Progression free survival (PFS) of patients treated with Sunitinib vs the combination of Axitinib + Avelumab d) Progression free survival (PFS) of patients treated with Sunitinib belonging to different angio-immune subtypes e) Progression free survival (PFS) of patients treated with combination of Axitinib + Avelumab belonging to different angio-immune subtypes f) Progression free survival (PFS) of patients belonging in C1 treated with Sunitinib vs the combination of Axitinib + Avelumab g) Progression free survival (PFS) of patients belonging in C2 treated with Sunitinib vs the combination of Axitinib + Avelumab h) Progression free survival (PFS) of patients belonging in C3 treated with Sunitinib vs the combination of Axitinib + Avelumab * = p<0.05, ** = p < 0.01, *** = p < 0.001, **** = p < 0.0001; One way ANOVA was used for comparison across more than two groups; Log rank tests were used for all survival analysis; error bars indicate SEM.

Median PFS improvement for patients with pre-treatment RNA sequencing data was from 8.4 months in the sunitinib arm to 12.5 months in the combination arm (Fig. 5c). We tracked survival across clusters to identify if the angio-immune subtypes can inform treatment choice in this setting. When treated with sunitinib, the angio-immune subtypes presented no significant differences in median PFS (Fig. 5d). However, when treated with the combination of avelumab and axitinib, the median PFS of C3 had not matured, C2 had a median PFS of 12.2 months, and C1 had a median PFS of 9.7 months (Fig. 5e). To determine if patients belonging to C1, C2, and C3 derive clinical benefit from the combination of avelumab and axitinib, PFS was tracked among the angio-immune subtypes across treatment arms. Remarkably, patients belonging to C1 derived no clinical benefit from the combination of avelumab and axitinib compared to sunitinib (Median PFS = 9.7 vs. 9.7; p = 0.668; Fig. 5f). Patients belonging in C2 also derived no clinical benefit from the combination (Median PFS = 12.2 months with combination treatment vs 8.3 months with sunitinib; p = 0.1469; Fig. 5g). In the C3 subtype, patients treated with avelumab and axitinib combination displayed significantly improved median PFS than the sunitinib arm (not matured vs. 8.2 months; p < 0.001; Fig. 5h). This analysis suggests that the angio-immune subtypes can be used to potentially exclude patients from treatment that do not derive clinical benefit but are faced with adverse toxicities from the therapy.

## DISCUSSION

ICB has revolutionized outcomes for a subset of cancer patients. However, a large proportion of patients fail to mount an effective and durable responses to ICB treatment. It is imperative to 1) identify strategies to improve response rates for patients upon ICB treatment and 2) identify pre-treatment characteristics of patients that can provide an a priori prediction to patients’ response to treatment. Such efforts can minimize toxicity profiles faced by patients while maximizing clinical benefits derived from ICB. Toward this effort, TME classification based on immunophenotype reported in few previous studies seems to predict the ICB therapy response^2, 23, 24, 25, 26, 27, 28^. These studies include the pan-cancer classification approaches such as immuno-phenoscore ^24^, the immuno-predictive score^25^, or the six TCGA immune TME subtypes^2^. PD-L1^26^, IFN*γ* ^27^, MHCI and II expression ^21, 28^, and CXCL9 expression ^29^ has also been reported to be a biomarker for the ICB response. However, these studies have restricted scopes, as they examined a subset of cancer types, and the prediction does not correlate well with the patient’s overall survival (OS) or progression-free survival (PFS). Recently, Bagaev et al.^6^ also analyzed the transcriptome of the TCGA to predict response to immunotherapy. In this work, the authors used functional gene expression signatures (Fges) representing the major functional components and immune, stromal, and other cellular populations of the tumor to classify TME into immune-enriched, fibrotic (IE/F), immune-enriched, non-fibrotic (IE), fibrotic (F), and immune-depleted (D). However, the fibrotic and immune scores do not seem to clearly predict the ICB response, suggesting the need to identify additional TME component that can correctly predict the ICB response.

Recent pre-clinical and clinical studies suggest that the status of tumor blood vessels can dictate immune responses in the TME. As such, characterizing the baseline angiogenic state and corresponding T-cell immune activity may provide us with tools to better inform treatment decision-making processes and delineate resistance mechanisms to ICB. Herein, we demonstrate that angiogenic activity and T-cell mediated immunity are inversely correlated across patients with 30 non-hematological solid tumor types. Distinct distribution of angiogenic and T-cell activity enrichment across tumor types enabled the stratification of patients into three conserved angio-immune subtypes. While tumor heterogeneity plagues efforts to develop overarching rules to define tumors independent of tissue of origin, the remarkable conservation of the distinct relationship between angiogenesis and T-cell mediated immunity across solid tumors enables us to develop highly interpretable rules to characterize tumors across tumor types.

The vessel normalization hypothesis provides a framework for the interaction of blood vessels and infiltrating immune cells in the TME ^30^. Highly angiogenic vascular networks impair the infiltration of immune cells as they express lower levels of selectins and adhesion molecules imperative for the trafficking of leukocytes to the tumor parenchyma^11^. Abnormal tumor blood vessels can also inhibit T-cell effector activity by expressing FAS ligand and inhibitory ligands ^31^. Accordingly, immune features in C3 angio-immune subtype include improved infiltration of anti-tumor immune cells, higher cytotoxicity activity of CD8+ T-cells, higher expression of co-stimulatory molecules, and higher expression of markers of T-cell exhaustion. The highly inflamed profile of the C3 subtype is partly due to the high mutational burden and neoantigen load in patients. Importantly, we identified distinct differences in mutational profiles of patients belonging to different angio-immune subtypes. The distinct mutations are actionable and may present distinct treatment modalities for patients harboring differing angio-immune subtypes.

ICB treatment allows the silencing of inhibitory signals on T-cells to enable reactivity against tumor cells. The current clinical landscape of treatment with ICB targets two key checkpoint molecules: PD1 and CTLA4. CTLA4 is an inhibitory receptor that competes with CD28 for B7 binding ^32^. B7 expression is primarily restricted to antigen-presenting cells in the tumor-draining lymph nodes ^33^. As such, the anatomical location of action for anti-CTLA4 therapeutics is primarily in lymph nodes. In comparison, anti-PD1/PDL1 treatments act in tumor cores. Tumor upregulation of PD-L1 engages with inhibitory PD1 receptors on T-cells ^34^. TME characteristics may have a more direct impact on anti-PD1/PDL1 efficacy. Local immune characteristics of patients belonging to the C3 subtype provide a compelling rationale for exploring treatment with anti-PD1/PDL1 ICB.

Patients with low angiogenesis levels and corresponding high T-cell activity belonging to the C3 subtype consistently respond better to ICB in melanoma, gastric cancer, bladder cancer, and renal cell cancer. Notably, the immune features that are characteristic of the C3 subtype alone fail to effectively predict responses to ICB in patients with renal, gastric, and bladder cancer^35^, suggesting the essential role of angiogenesis dictating the kind of immune response required to confer responses. The importance of the baseline angiogenic state provides a rationale for a temporal treatment strategy characterized by vascular normalization followed by ICB.

Re-evaluation of the Javelin Renal 101 clinical trial provides therapeutic relevance of the identified angio-immune subtypes. Patients harboring the C3 subtype demonstrated remarkable responses to the combination of axitinib and avelumab and received clinically significant improvements in survival compared to the previous standard of care sunitinib. Conversely, patients in the C2 and C1 subtypes did not benefit from combination of axitinib and avelumab compared to the previous standard of care sunitinib. This analysis suggests that we can potentially exclude a large subset of patients from the highly toxic combination treatment as they do not derive clinical benefit.

The angio-immune subtypes provide a more robust prognostic tool for ICB response and survival in comparison to previous studies. Importantly, we identify that the baseline angiogenic state can help dictate response to immune checkpoint blockade. Future investigations are required to explore accessible surrogates for local angiogenic and immune activity. Serum levels of vascular endothelial growth factor (VEGF) have been shown to predict the angiogenic capacity of non-small cell lung cancer ^36^. Whether this finding is consistent with other tumor types remains to be seen. Similarly, circulating CD8+ T cells can be probed for their cytolytic activity against tumors. Surrogate markers for tumor angiogenesis and T-cell activity would allow us to classify patients into angio-immune subtypes without a core biopsy. Additionally, the newly evolved vascular normalization hypothesis calls for a revisiting of dosage and combination schemes to optimize the use of anti-angiogenics in the clinic. High toxicity profile of the current anti-angiogenics mainly focusing on the VEGF pathway remains an obstacle. Recently uncovered anti-angiogenic targets like *MYCT1* that are dispensable for normal vessel maintenance provide early pre-clinical promise for a more tolerable anti-angiogenic therapeutic strategy ^11^. Such efforts will help better inform treatment decisions and enable us to maximize clinical benefit.

## METHODS

### TCGA Data

We used RNA, mutations, and clinical profiles for thirty non-hematological TCGA tumor types. Cancer types profiled include: Adrenocortical carcinoma (ACC), Bladder Urothelial Carcinoma (BLCA), Brain Lower Grade Glioma (LGG), Breast invasive carcinoma (BRCA), Colon adenocarcinoma (COAD), Cervical squamous cell carcinoma and endocervical adenocarcinoma (CESC), Cholangiocarcinoma (CHOL), Esophageal carcinoma (ESCA), Glioblastoma multiforme (GBM), Head and Neck squamous cell carcinoma (HNSC), Kidney Chromophobe (KICH), Kidney renal clear cell carcinoma (KIRC), Kidney renal papillary cell carcinoma (KIRP), Liver hepatocellular carcinoma (LIHC), Lung adenocarcinoma (LUAD), Lung squamous cell carcinoma (LUSC), Mesothelioma (MESO), Ovarian serous cystadenocarcinoma (OV), Pancreatic adenocarcinoma (PAAD), Pheochromocytoma and Paraganglioma (PCPG), Prostate adenocarcinoma (PRAD), Rectum adenocarcinoma (READ), Sarcoma (SARC), Skin Cutaneous Melanoma (SKCM), Stomach adenocarcinoma (STAD), Testicular Germ Cell Tumors (TGCT), Thyroid carcinoma (THCA), Uterine Carcinosarcoma (UCS), Uterine Corpus Endometrial Carcinoma (UCEC), and Uveal Melanoma (UVM).

#### a) RNA sequencing data

RNA sequencing data for 11,069 patients was downloaded from the GDC pan cancer portal (https://gdc.cancer.gov/about-data/publications/pancanatlas) ^37^. Data was processed using the Firehose pipeline with upper quantile normalization. For patients with more than one RNA-seq sample, primary tumor sample was favored. RNA sequencing samples from patients with DLBC and LAML were excluded.

#### b) Mutations

Version 2.8 of the mutations annotation file (MAF) generated by the MC3 group was downloaded from the GDC pan cancer portal (https://gdc.cancer.gov/about-data/publications/pancanatlas) ^37^. Samples from patients with DLBC and LAML were excluded. Cluster annotations were added to the MAF files. The maftools R package was used for visualization purposes.

#### c) Clinical data

Survival information was derived from TCGA-Clinical Data Resource (CDR) Outcome file provided in the GDC pan cancer portal (https://gdc.cancer.gov/about-data/publications/pancanatlas) ^37^.

### Patient Stratification

#### a) Building of the angio-immune score matrix

An unbiased selection of all gene sets relating to endothelial cell activity and T-cell activity from the molecular signatures database was conducted ^38^. A total of 91 gene signatures were identified and compiled to curate the angio-immune gene set collection. Gene set variation analysis (GSVA) was implemented to score the enrichment of 91 gene sets among patients of 30 TCGA cohorts to generate matrix of enrichment scores.

#### b) Clustering gene sets

Correlation matrix using Pearson coefficients were generated across enrichment scores for individual gene sets. Pheatmap package in R was used to generate heatmap visualization ^39^. Two modules of gene sets were identified and characterized based on the gene set membership.

#### c) Clustering patients

Correlation matrix using Pearson coefficients were generated across enrichment scores for patients. Pheatmap package in R was used to generate heatmap visualization. Three angio-immune subsets were identified and characterized based on unique distribution of enrichment of different gene set.

### Immune Characteristics of Tumors

#### a) Immune and Stromal Cell Enrichment

xCell, a gene signatures-based enrichment approach, was used to delineate enrichment of 64 immune and stromal cell types as previously described ^14^. Briefly, the xCell R package was used generate raw enrichment scores, transform into linear scale, and apply a spillover compensation to derive corrected enrichment scores. Distribution of enrichment scores for patients belonging to different angio-immune clusters were compared. Neoantigen load and TCR richness were downloaded from the GDC pan cancer portal (https://gdc.cancer.gov/about-data/publications/pancanatlas).

### Immunotherapy datasets

Pre-treatment TPM normalized RNA sequencing data from anti-PD1 treated cohorts were downloaded for the following studies: Gide ^18^, Hugo ^19^, Liu ^21^, Mariathasan ^20^, Kim ^22^, and Motzer ^40^. Studies were selected based on the following criteria: >20 samples for enrichment calculation, RNA sequencing of pre-treatment biopsies, and only anti-PD1/PDL1 treated patients. Gene signature enrichment and molecular subtypes were derived as described elsewhere in the manuscript. Survival and response rates to treatment when available were compared among angio-immune clusters.

### Statistics and Visualization

Unless indicated elsewhere, all visualizations were completed in GraphPad Prism 9. GraphPad Prism 9 was used for all statistical analysis. Data is presented as mean ± standard error of mean. One way ANOVA was used for comparison across more than two groups. Non-parametric tests were used when data was not normally distributed. Log rank tests were used for all survival analysis. Alpha value of 0.05 was used throughout the study.

## Acknowledgements

This work was supported by the NIH T32 AI007163 (D.A.G.B), the NIH grants R01HL149954 (K.C.), R01HL55337 (K.C.), and Siteman Investment Program Research Development Awards (K.C.).

